# Acid excretion is impaired in calcium oxalate stone formers

**DOI:** 10.1101/2024.09.30.615797

**Authors:** Pedro H Imenez Silva, Nasser A. Dhayat, Daniel G. Fuster, Harald Seeger, Alexander Ritter, Thomas Ernandez, Florian Buchkremer, Beat Roth, Olivier Bonny, Isabel Rubio-Aliaga, Carsten A. Wagner

## Abstract

**Background:** Urine pH is a key factor in kidney stone formation. We aimed to identify whether acid excretion capacity is disturbed in calcium oxalate (CaOx) or calcium phosphate (CaP) stone formers.

**Method:** Urinary, serum, clinical, and anthropomorphic baseline data were obtained from the Swiss Kidney Stone Cohort, a prospective, longitudinal, and multi-centric observational study. We included in this study 193 non-stone formers (NSF, confirmed by negative CT scan), and 309 CaOx and 28 CaP stone formers. Titratable acids, net acid excretion (NAE), NAE capacity (NAEC) and acid-base (AB) score were calculated. Logistic regression analyses were used to estimate the potential associations of various acid-base variables with the occurrence of CaOx kidney stones.

**Results:** CaOx stone formers showed a disturbed capacity to excrete acids in comparison to NSF (NAEC NSF = 3.49±12.6 mmol/24h; CaOx = −1.06±13.10; CaP = 0.97±14.70 and AB score NSF = 20.5±6.36 mmol/24h; CaOx = 17.9± 6.53; CaP = 18.8±6.10). The correlation between urine calcium and urine pH was altered in CaOx stone formers and between urine calcium and NAE was stronger in CaP stone formers. Logistic models showed that urinary ammonium was negatively associated with CaOx stone formation (unadjusted model, odds ratio 0.43[0.32-0.58], p< 0.001 for CaOx). Urine calcium was positively associated with CaOx kidney stones (2.85 [2.11-3.92], p<0.001). Similar results were obtained after adjusting for age, sex, and BMI. Replacing urine ammonium, pH, and phosphate with NAEC or ammonium and pH with AB score in our logistic regression models showed that NAEC and AB score are strongly associated with CaOx kidney stone formation.

**Conclusion:** Ammonium excretion, NAEC and AB score are associated with the occurrence of CaOx kidney stones suggesting a potential role of proximal tubule dysfunction in their formation. CaP stone formers exhibit a disproportionately higher calcium excretion when acid excretion increases.

**Key learning points:** *What was known:* Urine pH is a strong determinant in the formation of various urologically relevant crystals. Impaired urine acidification capacity has been observed in individuals who form calcium phosphate and uric acid stones.

*This study adds:* When compared to non-stone formers, calcium oxalate stone formers are marked by a reduced capacity of excreting acids when urine pH becomes more acidic.

*Potential impact:* The calculation of net acid excretion capacity and acid-base score are novel tools to identify those under potential higher risk of developing calcium oxalate stones.

## INTRODUCTION

Kidney stone disease has a high prevalence worldwide, leading to significant morbidity and potentially increasing the risk of chronic kidney disease [1, 2]. Dietary, lifestyle, genetic, and environmental factors are associated with higher risk of stone formation [3–5]. Yet, urine pH and supersaturation are the strongest factors underlying urine crystal formation [6]. The influence of pH on nephrolithiasis is further shown by the larger risk of kidney stone formation in patients with urine acidification defects, with this risk being reduced by alkali therapy (except in patients with urate stones) [7].

Acid-base status also influences the content of citrate in the urine, which impacts on the formation of calcium-containing kidney stones [8]. Metabolic acidosis and acidic urine pH increase citrate reabsorption in the kidney proximal tubules, while alkaline urine pH reduces citrate reabsorption [9]. However, how the renal capacity to excrete acid influences the incidence of common kidney stone types is not completely understood. Worcester et al. suggested that calcium phosphate (CaP) stone formers (SF) have a proximal tubule acidification defect while female CaOx SF have reduced intestinal alkali absorption, which could generate the acidic urine pH typically observed in this group (3).

Leveraging the extensive urine biochemistry profile available in the Swiss Kidney Stone Cohort, we tested whether urinary acid-base parameters, such as pH, ammonium, titratable acids, net acid excretion (NAE), and citrate were associated with CaOx and CaP stone formation. We assessed the kidney’s capacity of excreting acids by calculating net acid excretion capacity from the residuals of the relation between net acid excretion and urine pH and the recently published acid-base (AB) score [10].

## METHODS

### Swiss Kidney Stone cohort

The Swiss Kidney Stone cohort (SKSC, ClinicalTrials.gov, NCT01990027) is an investigator-initiated prospective, multicentric longitudinal, observational study in patients with kidney stones. The study is performed according to the current version of the Declaration of Helsinki, ICH-GCP, GEP, and Swiss law on human studies, and was approved by the Swiss Cantonal Ethics Committees [11]. Individuals with kidney stone disease (SF) were followed over 3 years to collect blood and urine samples and clinical data. This study analyzed baseline data collected at visit 2 (V2), which was performed 2 weeks after the screening visit. Patients had their last stone episode between 6 and 15 weeks prior to V2. A control cohort of individuals CT-proven free of kidney stones and calcifications has been recruited (NSF, n = 205).

SKSC included individuals with recurrent stone disease or specified risk factors [11]. The exclusion criteria for stone formers in SKSC were: age below 18 years old, no signed informed consent and inability to follow the study protocol. Here, we included only SF with stones classified as CaOx (n = 333) or CaP (n = 135). CaOx SF were defined as those with an average stone mineral composition consisting of >50% calcium oxalate. Patients with >50% CaP content were categorized as CaP stone formers. We excluded individuals with missing data on kidney stone type (n = 165), or urine pH, citrate, phosphate, magnesium, or oxalate (n = 125 SF and 11 NSF). We noticed that calculated bicarbonate and NAE were highly influenced by urine pH values above 7.5, which skewed the regression and correlation analyses. Therefore, 4 SF and 1 NSF with urine pH values above 7.5 were excluded from the analyses. Additional analyses were performed by excluding individuals who used supplements or medication with known effects on acid-base balance (Supplemental Table 1).

### Blood and urine data

In this study we only used urine collected under mineral oil (paraffin) containing thymol as a chemical preservative. 24h urine samples were analyzed for urine volume, Na^+^, K^+^, Cl^−^, Ca^2+^, inorganic phosphate (Pi), Mg^2+^, total protein, albumin, oxalate, citrate, creatinine, uric acid, urea, ammonium, pH, and sulfate. Blood was analyzed for Na^+^, K^+^, Cl^−^, Ca^2+^, Mg^2+^, albumin, phosphate, glucose, creatinine, urea, uric acid, parathyroid hormone, calcidiol (25-OH-vitamin D^3^), calcitriol (1,25-OH2-vitamin D^3^), intact FGF23, HbA1c, and lipids. A minor fraction of patients also had blood pH (n=148) assessed.

### Equations

Estimated glomerular filtration rate (eGFR) was calculated using the CKD-EPI 2009 formula without the race coefficient [12]. NAEC was calculated from the residuals of the linear regression between urine pH and ammonium excretion as done in [13]. AB score was calculated as described in [10] (Table 1). Other equations were calculated similar to Ferraro et al [14] (Table 1).

**Table 1:**
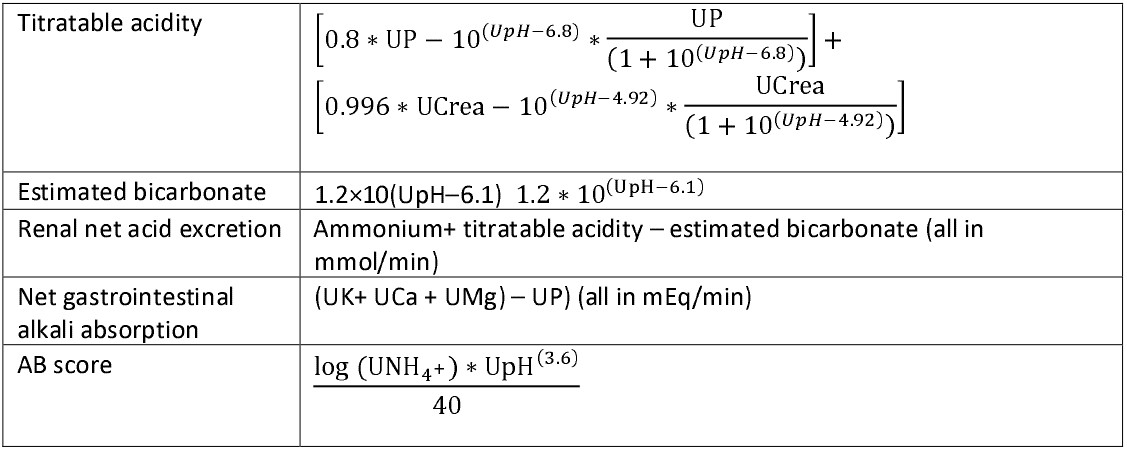
Equations used in this study. U before an abbreviation indicates Urine. Crea – creatinine, P– phosphorus, K – potassium Ca –calcium, Mg – magnesium.

### Statistical Analysis

Logistic regression models were performed with centralized data normalized to their standard deviations. Linearity between continuous covariates and the dependent variable was assessed via a component residual plot. Multicollinearity was evaluated by calculating variance inflation factor (VIF). Variables used in the logistic models were considered linear and without multicollinearity.

Pearson correlation coefficients were obtained for correlation analysis. To assess whether the correlation results are influenced by kidney stone type as a potential effect modifier, additional linear regression models were performed, incorporating kidney stone type as an interaction term.

Continuous data are shown as mean ± SD and skewed data as median + [25th percentile; 75th percentile]. Categorical data are shown as count number and % within-group. Odds ratio (OR) are shown as OR [95% confidence interval]. Multi-group comparisons were performed with ANOVA followed by Bonferroni correction. The α used for all analyses was 0.05.

All data organization, cleaning, and transformation, and calculations of descriptive statistics, statistical analyses, and visualization were performed on the R environment [15] with Rstudio and using the following packages: car, dplyr, ggplot2, ggpubr, Hmisc, janitor, lubridate, rstatix, summarytools, table1.

## RESULTS

We included 309 calcium oxalate (CaOx) and 28 calcium phosphate (CaP) stone formers (SF) and 193 non-stone formers (NSF) in this study (Figure 1). CaOx SF were slightly older than NSF individuals and CaP SF. They were more often males, and had slightly higher BMI than NSF, besides similar baseline eGFR and showed lower calculated gastrointestinal alkali absorption levels (Table 2). Baseline blood and urinary parameters are summarized in Supplemental Tables 2 and Table 3. 24h urine excretion in CaOx and CaP SF was lower for citrate, oxalate, and potassium (Table 3), and in addition, CaOx SF exhibited a lower 24h urine volume, ammonium and magnesium excretion. In contrast, CaOx and CaP SF excreted more calcium (Table 3). Urine pH was more acidic in CaOx SF in comparison with NSFs.

**Table 2.** Baseline characteristics. Categorical data are reported as count and percentage in relation to each group’s total sample size and continuous data as mean ± SD. Parameters with missing data have their sample size (n) shown next to the mean or median value of each group. GI = gastrointestinal.

| Baseline characteristics |  |  |  |
| --- | --- | --- | --- |
|  | NSF<br>(N=193) | CaOx<br>(N=309) | CaP<br>(N=28) |
| Sex (Female) | 84 (43.5%) | 89 (28.8%) | 18 (64.3%) |
| Age (years) | 43.0 ± 13.6 | 46.4 ± 14.3 | 44.3 ± 14.4 |
| BMI (kg/m<sup>2</sup>) | 25.0 ± 4.12 | 26.9 ± 4.66 | 26.4 ± 5.73 |
| Weight (kg) | 75.7 ± 15.1 | 80.7 ± 16.4 | 74.7 ± 21.2 |
| eGFR by CKD-EPI<br>(ml/min per 1.73m<sup>2</sup>) | 97.3 ± 16.1 | 96.8 ± 18.5 | 98.9 ± 16.8 |
| GI alkali absorption<br>(mEq/min) | 29.6 ± 23.7 | 20.8 ± 15.6 | 21.0 ± 11.8 |
| Hypertension | 18 (9.4%), n =192 | 69 (22.3%), n = 307 | 9 (32.1%) |
| Diabetes type 1 | 0 (0%), n = 192 | 1 (0.3%), n = 305 | 0 (0.0%) |
| type 2 | 3 (1.6%), n = 192 | 28 (9.1%), 305 | 2 (7.1%) |
| Dyslipidemia | 22 (11.5%), n = 192 | 41 (13.3%), n = 258 | 1 (3.6%), n = 25 |
| Gout | 2 (1.0%), n = 192 | 6 (1.9%), n = 305 | 1 (3.6%) |
| Smoker | 43 (22.4%), n = 190 | 85 (27.6%), n = 301 | 5 (17.9%) |
| Ex-smoker | 26 (13.5%), n = 190 | 38 (12.3%), n = 301 | 2 (7.1%) |
| Osteoporosis | 5 (2.6%), n = 189 | 8 (2.6%), n = 232 | 0 (0.0%), n = 22 |
| Fractures | 96 (50.0%), n = 191 | 75 (24.3%), n = 296 | 4 (14.3%), n = 26 |
| Immobilization | 14 (7.3%), n = 180 | 15 (4.9%), n = 249 | 3 (10.7%), n = 23 |
| Kidney stones in the family | 29 (15.1%), n = 187 | 144 (46.4%), n = 305 | 14 (50.0%), n = 27 |
| Kidney transplantation | 1 (0.5%), n = 185 | 0 (0%), n = 302 | 1 (3.6%) |
| Pyelonephritis in the past | 7 (3.6%) | 48 (15.5%) | 11 (39.3%) |
| Nephrocalcinosis | 0 (0%), n = 191 | 19 (6.1%), n = 256 | 3 (10.7%), n = 25 |
| Inflammatory bowel diseases | 2 (1.0%), n = 189 | 17 (5.5%), n = 304 | 2 (7.1%) |
| Gastric bypass | 2 (1.0%), n = 190 | 20 (6.5%), n = 304 | 0 (0.0%) |
| Vegetarian | 16 (8.3%) | 10 (3.2%) | 1 (3.6%) |
| Lactose-free | 9 (4.7%) | 8 (2.6%) | 0 (0%) |
| Vegan | 2 (1.0%) | 0 (0%) | 0 (0%) |
| Other diets | 15 (7.8%) | 10 (3.2%) | 3 (10.7%) |
| Supplements and vitamins | 64 (33.3%), n=188 | 67 (21.7%), n = 250 | 8 (28.6%), n = 21 |

**Table 3.**
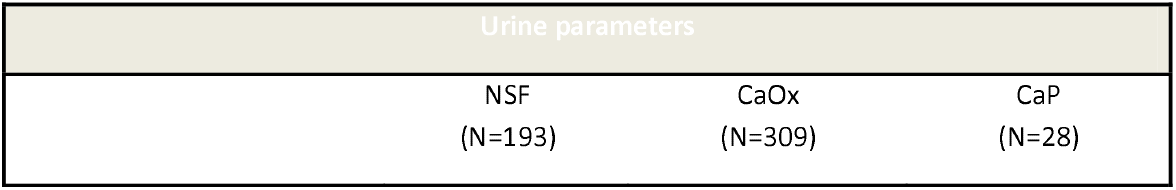
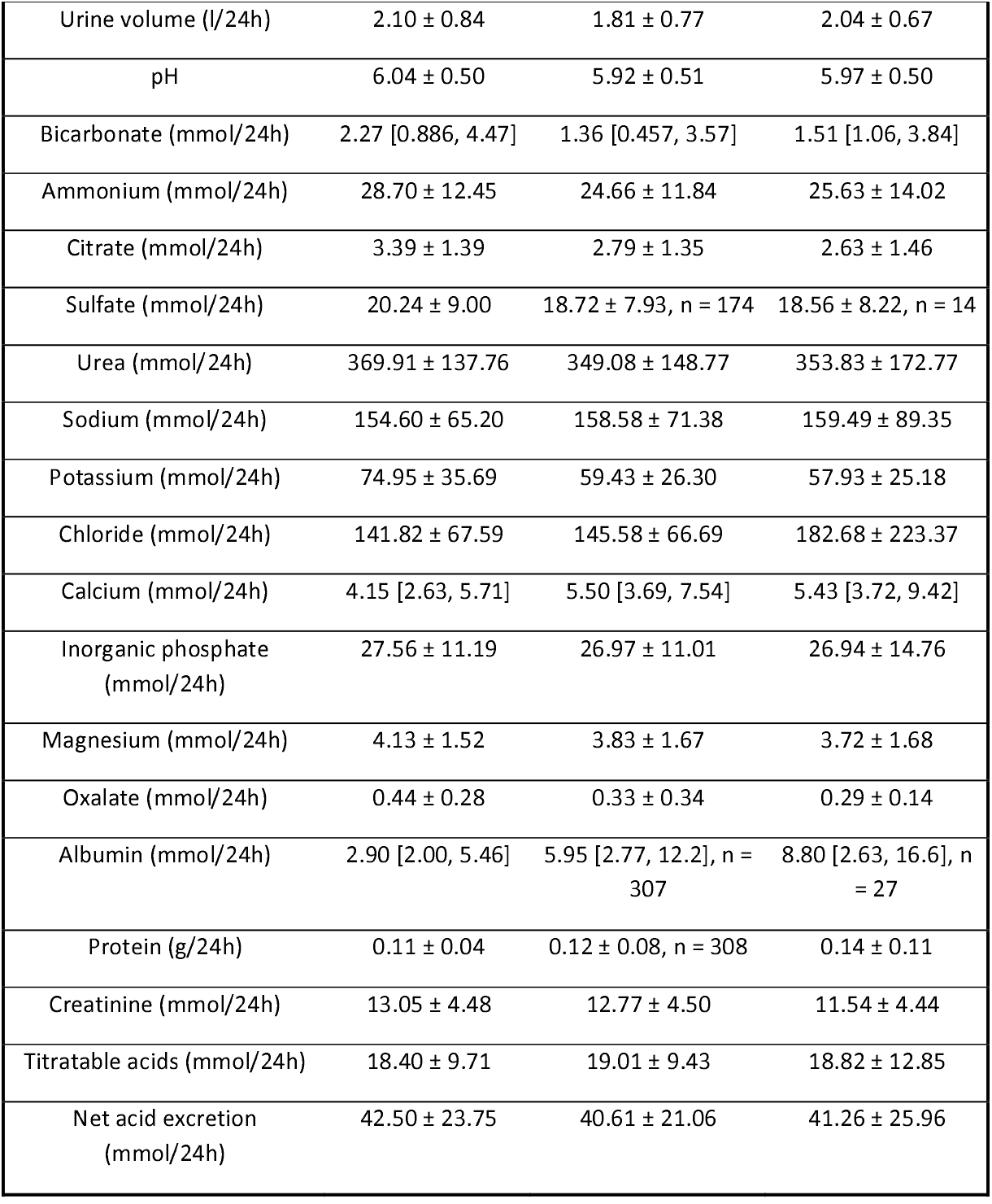
Urine parameters in SF and NSF individuals. Baseline urine data are shown as mean ± SD and continuous skewed data as median [25th percentile; 75th percentile]. Parameters with missing data have their sample size (n) shown next to the mean or median value of each group. NAE = net acid excretion

| Urine parameters |  |  |
| --- | --- | --- |
| NSF<br>(N=193) | CaOx<br>(N=309) | CaP<br>(N=28) |

Table 3. Urine parameters in SF and NSF individuals.
|  |  |  |  |
| --- | --- | --- | --- |
| Urine volume (l/24h) | 2.10 ± 0.84 | 1.81 ± 0.77 | 2.04 ± 0.67 |
| pH | 6.04 ± 0.50 | 5.92 ± 0.51 | 5.97 ± 0.50 |
| Bicarbonate (mmol/24h) | 2.27 [0.886, 4.47] | 1.36 [0.457, 3.57] | 1.51 [1.06, 3.84] |
| Ammonium (mmol/24h) | 28.70 ± 12.45 | 24.66 ± 11.84 | 25.63 ± 14.02 |
| Citrate (mmol/24h) | 3.39 ± 1.39 | 2.79 ± 1.35 | 2.63 ± 1.46 |
| Sulfate (mmol/24h) | 20.24 ± 9.00 | 18.72 ± 7.93, n = 174 | 18.56 ± 8.22, n = 14 |
| Urea (mmol/24h) | 369.91 ± 137.76 | 349.08 ± 148.77 | 353.83 ± 172.77 |
| Sodium (mmol/24h) | 154.60 ± 65.20 | 158.58 ± 71.38 | 159.49 ± 89.35 |
| Potassium (mmol/24h) | 74.95 ± 35.69 | 59.43 ± 26.30 | 57.93 ± 25.18 |
| Chloride (mmol/24h) | 141.82 ± 67.59 | 145.58 ± 66.69 | 182.68 ± 223.37 |
| Calcium (mmol/24h) | 4.15 [2.63, 5.71] | 5.50 [3.69, 7.54] | 5.43 [3.72, 9.42] |
| Inorganic phosphate (mmol/24h) | 27.56 ± 11.19 | 26.97 ± 11.01 | 26.94 ± 14.76 |
| Magnesium (mmol/24h) | 4.13 ± 1.52 | 3.83 ± 1.67 | 3.72 ± 1.68 |
| Oxalate (mmol/24h) | 0.44 ± 0.28 | 0.33 ± 0.34 | 0.29 ± 0.14 |
| Albumin (mmol/24h) | 2.90 [2.00, 5.46] | 5.95 [2.77, 12.2], n = 307 | 8.80 [2.63, 16.6], n = 27 |
| Protein (g/24h) | 0.11 ± 0.04 | 0.12 ± 0.08, n = 308 | 0.14 ± 0.11 |
| Creatinine (mmol/24h) | 13.05 ± 4.48 | 12.77 ± 4.50 | 11.54 ± 4.44 |
| Titrateable acids (mmol/24h) | 18.40 ± 9.71 | 19.01 ± 9.43 | 18.82 ± 12.85 |
| Net acid excretion (mmol/24h) | 42.50 ± 23.75 | 40.61 ± 21.06 | 41.26 ± 25.96 |

**Figure 1:**
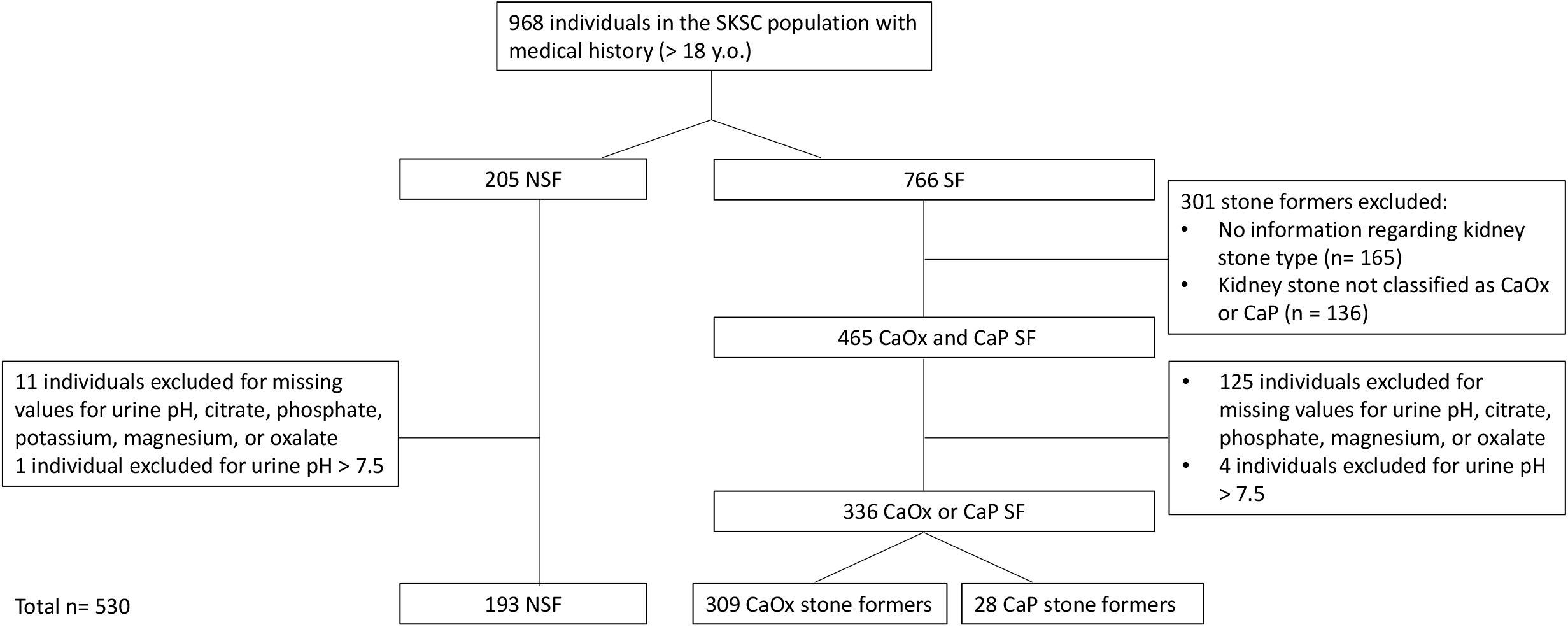
Flow chart of the studied population derived from the Swiss Kidney Stone Cohort. CaOx = calcium oxalate, CaP = calcium phosphate, and NSF = non-stone former.

Next, we tested whether urinary acid-base parameters were associated with the presence of CaOx or CaP stones in our patients. In addition, we included variables typically associated with kidney stone formation, such as urinary calcium and oxalate excretion. Urine magnesium is a relevant mediator [16] or confounder [17] in this model and was also included. We performed logistic regression models using urine ammonium, pH, citrate, phosphate, calcium, magnesium, and oxalate as independent variables (model 0). We also corrected this model using age, BMI, and sex as confounders given their potential effects both on exposure and outcome (model 1). According to model 0 for CaOx, holding all other independent variables constant, urine ammonium, pH, citrate, and calcium were significantly associated with the odds of having kidney stones (Table 4). Results remained largely consistent after adjusting for age, BMI, and sex, although urine pH appeared to have been partially reduced in effect size (Supplemental Table 3). Urinary ammonium had the lowest and urinary calcium the highest odds ratio for the association with CaOx occurrence in the adjusted logistic regression analysis. Given the low number of CaP SF in our cohort, logistic models did not perform well and data are not included.

**Table 4.** Logistic regression model 0 with CaOx stone formation as dependent variable. OR = odds ratio. The p-values were obtained from a Wald test with α = 0.05.

| Urine parameter | estimate | Std error | statistic | p-value | OR | 2.50% | 97.50% |
| --- | --- | --- | --- | --- | --- | --- | --- |
| (Intercept) | 0.60 | 0.11 | 5.599 | <0.001 | 1.83 | 1.48 | 2.27 |
| Ammonium | -0.84 | 0.16 | -5.403 | <0.001 | 0.43 | 0.32 | 0.58 |
| pH | -0.37 | 0.11 | -3.297 | 0.001 | 0.69 | 0.56 | 0.86 |
| Citrate | -0.52 | 0.12 | -4.373 | <0.001 | 0.60 | 0.47 | 0.75 |
| Phosphate | 0.23 | 0.15 | 1.51 | 0.131 | 1.26 | 0.93 | 1.72 |
| Calcium | 1.05 | 0.16 | 6.654 | <0.001 | 2.85 | 2.11 | 3.92 |
| Magnesium | -0.28 | 0.15 | -1.882 | 0.060 | 0.76 | 0.57 | 1.01 |
| Oxalate | -0.19 | 0.15 | -1.275 | 0.202 | 0.82 | 0.60 | 1.10 |

Urine acid-base parameters were differentially affected in CaOx and CaP SF. We calculated net acid excretion capacity (NAEC) from the residuals of the regression analysis between NAE and urine pH [13] (Figure 2A). Considering that NAE and urine pH are strongly associated in healthy kidneys, NAEC provides an index of how much acid is excreted for a given urine pH. NAEC is reduced in CaOx SF (−1.74 ± 17.78) in relation to NSF (3.80 ± 17.87, p = 0.002). CaP SF showed values that were intermediate between the other two groups (0.63 ± 20.84, p = 1.0) (Figure 2B). Calculated titratable acidity was similar across all groups (Figure 2C). Recently, another approach has been proposed to analyze the relation between ammonium excretion and urine pH and was termed acid-base score (AB score) [10]. AB score was reduced in CaOx SF in comparison with NSF (20.37 ± 7.08 mmol/24h vs 23.14 ± 6.97, p < 0.001) (Figure 2D). Both NAEC and AB score demonstrate that CaOx SF have an acid excretion disorder.

**Figure 2:**
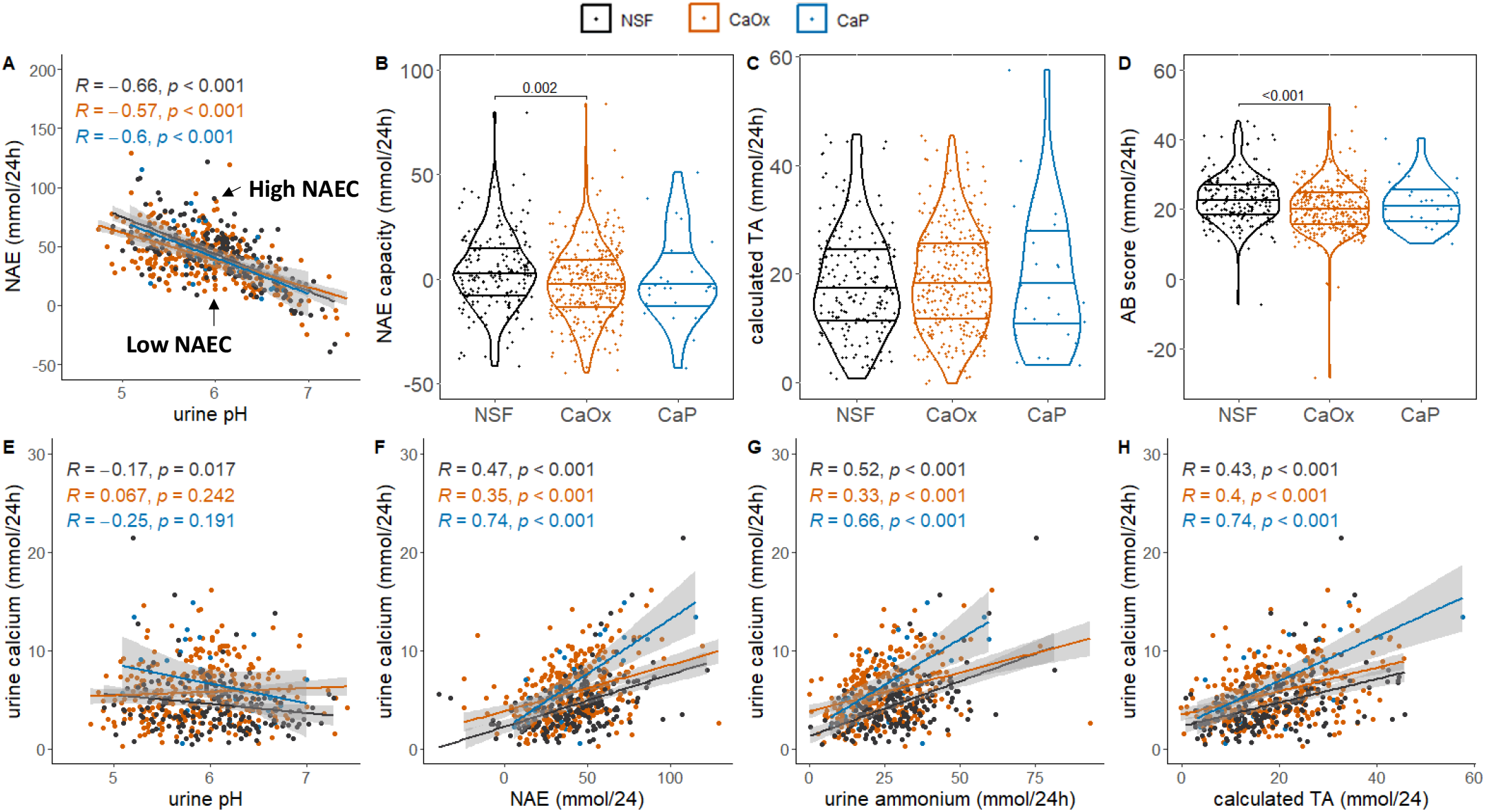
Acid excretion capacity is disturbed in CaOx stone formers (SF) and acid-dependent calcium excretion is exacerbated in CaP SF. (A) Pearson correlation analysis between urine pH and net acid excretion (NAE). (B-D) Violin plots showing median and 25th and 75th percentile of (B) NAE capacity, (C) calculated titratable acidity (TA), and (D) acid-base (AB) score. (E-H) Pearson correlation analysis between urine pH and urine calcium (E), NAE and urine calcium (F), urine ammonium and calcium (G), and calculated TA and urine calcium (H). Blue dots = non stone formers (NSF), orange dots = CaOx SF, black dots = CaP SF. R = Pearson correlation coefficient and p = p-value associated with this correlation. α = 0.05.

Next, we performed logistic regression analysis for CaOx stone occurrence replacing urine ammonium, pH, and phosphate with NAEC. NAEC was associated with CaOx stone occurrence (OR 0.55 [0.42-0.71], p < 0.001) (Table 5). AB score also associated with CaOx stone occurrence (OR 0.67 [0.53-0.83], p < 0.001) in a model replacing urine ammonium and pH with AB score (Table 6). NAEC and AB score remained associated with CaOx occurrence after adjustment for age, BMI, and sex. While the odds ratio for NAEC were further reduced (OR 0.48 [0.36-0.64], p < 0.001), AB score showed a weaker effect (OR 0.73 [0.58-0.93], p = 0.011) (Supplemental Tables 4 and 5).

**Table 5.** Logistic regression model 0 with CaOx stone formation as dependent variable and NAEC replacing ammonium, pH, and phosphate. OR = odds ratio. The p-values were obtained from a Wald test with α = 0.05.

| Urine parameter | estimate | Std error | statistic | p-value | OR | 2.50% | 97.50% |
| --- | --- | --- | --- | --- | --- | --- | --- |
| (Intercept) | 0.59 | 0.11 | 5.606 | <0.001 | 1.81 | 1.48 | 2.24 |
| NAEC | -0.60 | 0.13 | -4.531 | <0.001 | 0.55 | 0.42 | 0.71 |
| Citrate | -0.47 | 0.11 | -4.167 | <0.001 | 0.62 | 0.49 | 0.77 |
| Calcium | 1.08 | 0.15 | 6.953 | <0.001 | 2.93 | 2.19 | 4.01 |
| Magnesium | -0.21 | 0.14 | -1.549 | 0.122 | 0.81 | 0.61 | 1.06 |
| Oxalate | -0.27 | 0.15 | -1.836 | 0.066 | 0.76 | 0.56 | 1.01 |

**Table 6.** Logistic regression model 0 with CaOx stone formation as dependent variable and AB score replacing urine ammonium and pH. OR = odds ratio. The p-values were obtained from a Wald test with α = 0.05.

| Urine parameter | estimate | Std error | statistic | p-value | OR | 2.50% | 97.50% |
| --- | --- | --- | --- | --- | --- | --- | --- |
| (Intercept) | 0.59 | 0.10 | 5.601 | <0.001 | 1.80 | 1.47 | 2.21 |
| AB score | -0.40 | 0.11 | -3.569 | <0.001 | 0.67 | 0.53 | 0.83 |
| Citrate | -0.45 | 0.11 | -3.967 | <0.001 | 0.64 | 0.51 | 0.79 |
| Phosphate | -0.05 | 0.14 | -0.348 | 0.728 | 0.95 | 0.73 | 1.25 |
| Calcium | 0.95 | 0.15 | 6.219 | <0.001 | 2.59 | 1.93 | 3.53 |
| Magnesium | -0.32 | 0.14 | -2.227 | 0.026 | 0.73 | 0.55 | 0.96 |
| Oxalate | -0.31 | 0.15 | -2.097 | 0.036 | 0.73 | 0.54 | 0.97 |

Healthy kidneys increase the excretion of calcium in response to an acid load and calcium has been reported to be associated with urine pH and NAE [18, 19]. We performed Pearson correlation analysis to detect whether these correlations were preserved in stone formers. We identified a dissipation of these associations in CaOx SF (Figure 2E). The correlation between calcium excretion and urine pH was not strong but still significant in NSF individuals (R = −0.17, p-value = 0.017) and it was absent in CaOx SF (R= −0.07, p = 0.24) and CaP SF (R= −0.25, p = 0.19). Urine calcium was correlated with urine NAE, ammonium, and calculated TA in all three groups, but these correlations were stronger and steeper in CaP SF (Figure 2F-H) as shown by higher Pearson correlation coefficients and steeper slopes (e.g., slope of NAE vs urine calcium: 0.11 vs 0.05, p = 0.005 for interaction with CaP stone formation).

Next, we tested whether differences in intestinal alkali absorption could explain the abnormal excretion patterns for calcium and ammonium in calcium SF. Although intestinal alkali absorption was reduced in CaOx and CaP SF (Table 2), its relation with urinary acid-base parameters and urine calcium was not significantly altered in SF (Supplemental figure 1).

We repeated the Pearson correlation analysis shown in Figure 2 but removing data from individuals who reported the use of supplements or medications with known effects on acid-base balance (Supplemental Figures 2 and 3). Acid excretion capacity was similarly altered in this subpopulation when compared to individuals not excluded by supplement or medication intake. Moreover, we repeated the logistic regression analyses with this subpopulation. This analysis provided similar patterns to those reported in Tables 4-6, with ammonium and NAEC being strongly negatively associated with CaOx stones. Interestingly, AB score showed a weaker association with CaOx stone occurrence in this subpopulation (Supplemental Tables 10 and 11). Calcium remained positively associated with CaOx (Supplemental Tables 6-11). Therefore, the similar results before and after applying this exclusion criterion suggest that the pathomechanisms underlying calcium stone formation reported here are likely intrinsic to the tubular function of these patients.

## DISCUSSION

The main chemical determinants of urinary crystal formation are supersaturation, pH, and impurities [6]. Moreover, pH modulates different steps of crystal formation, such as supersaturation itself and nucleation [20]. While the contribution of urine pH to CaOx formation [21] is still unclear, CaP supersaturation increases abruptly between pH 6 and 7 [22] and an alkaline urine is one of the strongest determinants of CaP formation [8, 23]. In our cohort, CaOx SF showed a more acidic urine than NSF, but no difference was observed in CaP SF. Despite this alteration, urine pH was only weakly associated with CaOx formation in our logistic regression models, in agreement with another recent study (22). In contrast, other key acid-base variables, such as urine ammonium and citrate were strongly associated with CaOx kidney stones. Furthermore, CaOx SF showed reduced NAEC and AB scores, demonstrating a reduced acid excretion capacity in this population.

### Lower acid excretion capacity associates with CaOx kidney stones

Impaired acid-base handling has been previously suggested as a potential contributor to calcium stone formation [24]. Worcester et al compared urine composition of CaOx and CaP SF in fasting and fed conditions. They observed that CaP SF in fed conditions had higher ammonium but lower citrate excretion accompanied by higher urine pH and suggested that this would derive from a proximal tubule defect. In our study, samples were collected over 24 hours while participants were eating their normal diet and ammonium excretion was lower in CaOx SF. Under the same condition, Worcester et al reported no significant differences.

Previously, we showed that mice lacking Gpr68, the proton-activated ovarian cancer G protein-coupled receptor 1 (OGR1), have a proton secretion defect in the proximal tubule [25]. These animals show a strong disruption of the associations between urine pH and urinary excretion of ammonium and calcium [19]. Interestingly, when oxalate nephropathy was induced with a high oxalate diet, these animals deposited more calcium oxalate crystals and tubular damage [26]. While the present work does not demonstrate causality, we hypothesize that a reduced capacity to excrete acid may be linked to a higher CaOx crystal deposition.

### Origin of the defect in CaOx stone formers

Worcester et al suggested that a more acidic urine pH in fed female CaOx SF would derive from lower gastrointestinal alkali absorption [24]. Likewise, we identified reduced intestinal alkali absorption in CaOx and CaP SF, suggesting that dietary factors might be involved in the acid-base disturbances observed in calcium stone formers. However, no differences were found among all groups in the association between GI alkali absorption and net acid or ammonium excretion, or urine pH. Alterations in ammonium and net acid excretion capacity suggest that kidneys are the site responsible for the observed defects. It is tempting to speculate that the proximal tubule is the segment responsible for this defect, based on the following considerations: 1) ammoniagenesis occurs in the proximal tubules, and while its excretion depends on processes downstream in the nephron, they ultimately depend on the capacity to acidify urine [27], which is apparently preserved in CaOx stone formers; 2) a proximal tubule defect in acid secretion is enough to generate a higher risk for CaOx crystal deposition [19, 25, 26]; 3) In this study, CaOx stone formers showed also reduced urine citrate excretion and the proximal tubule is the main site of citrate reabsorption.

Lower ammonium excretion was positively associated with CaOx kidney stone types. While NAE did not associate with CaOx stones occurrence (data not shown), urine pH showed a weak association with it after adjustment for age, BMI, and sex; NAEC, being the residuals of the linear regression of NAE and urine pH, offers information that pH, NAE, and ammonium excretion cannot provide individually. The defect in ammonium excretion is further illustrated by the lower AB score, which has also been shown to be associated with a higher risk for CKD progression [10]. It remains to be determined whether individuals with CKD and low AB score have a higher risk for CaOx kidney stone deposition.

The low sample size for CaP stone formers limited us to apply the same logistic analysis for this population. However, mean levels for urine ammonium and citrate were similar between CaOx and CaP stone formers. To further test whether NAE capacity is altered in calcium stone formers, NAE capacity should be assessed in individuals challenged with an acid load. A few studies have subjected stone formers to an acid load, either via NH_4_Cl intake or furosemide with fludrocortisone [28–30]. Incomplete renal tubular acidosis was identified in a minor proportion of CaP stone formers and probably cannot explain most of the cases [28, 31].

### Exacerbated calcium excretion in CaP stone formers and dysregulated calcium excretion in CaOx stone formers

Hypercalciuria is a frequently observed feature in calcium stone formers [32]. Urine acidification increases calcium excretion and vice-versa [33–35]. In our study, we detected a weak negative association between urine pH and calcium excretion and a moderate positive association between NAE and calcium excretion in NSF. However, CaP SF exacerbated these interactions, in which calcium excretion was proportionally higher towards higher levels of acid excretion. These individuals showed both a higher Pearson correlation coefficient and a steeper slope when correlating both variables. CaOx SF showed an apparent disruption of the association between acid and calcium excretion despite also presenting higher calcium excretion levels in comparison with NSF. These results further demonstrate that calcium excretion is closely linked with acid excretion and that CaOx and CaP SF exhibit distinct dysregulations in the interactions between kidney calcium and acid handling.

Our study has some limitations. The low sample size for CaP stone formers may have prevented the detection of relevant associations between CaP stone formation and acid-base parameters. Titratable acidity and bicarbonate were calculated instead of measured, besides the worse performance of bicarbonate calculation when urine pH is high (i.e. above 7.5). However, in this study, calculated TA has an advantage over measured TA, as it is not affected by the precipitation of calcium phosphate [36]. Moreover, we did not include systemic acid-base status in our analysis given incomplete blood gas analysis data. We also did not stratify our cohort for sex to preserve statistical power. Yet, we adjusted all analyses for sex, which did not change outcomes. However, our study shows important strengths: extensive urinary biochemical data performed close to stone diagnosis, the validation of our control group by CT scan, and detailed medication history of patients.

In summary, the kidney’s capacity of excreting acid and its interaction with calcium excretion are differentially impaired in CaP and CaOx stone formers, which most probably reflects different tubular defects that increase the probability of stone formation.

## Acknowledgements

The Swiss Kidney Stone Cohort has been supported by the National Center of Competence in Research NCCR.CH financed by the Swiss National Science Foundation and the University of Zurich. CAW is also supported by the Swiss National Science Foundation (212303). DGF was supported by the Swiss National Science Foundation (grants # 33IC30-166785/1 and 320030-227578).

## Figure Legends

Supplemental Figure 1: Variation in gastrointestinal alkali absorption does not explain altered ammonium or calcium excretion. Correlation between calculated gastrointestinal alkali and urine pH (A), ammonium (B), calcium (C), and net acid excretion is not significantly modified in CaOx or CaP stone formers. Black dots = NSF, orange dots = CaOx CaOx stone formers, and blue dots = CaP stone formers. R = Pearson correlation coefficient and p = p-value associated with this correlation. α = 0.05.

Supplemental Figure 2: Number of individuals per group after exclusion by drugs and supplements that affect the acid-base status. ACEi = Angiotensin-converting enzyme inhibitors, ARBs = Angiotensin receptor blockers, IS = immunosuppressants.

Supplemental Figure 3: Intake of medication and supplements interacting with acid-base balance did not play a major role in acid excretion capacity and its relationship with calcium excretion. (A) Pearson correlation analysis between urine pH and net acid excretion (NAE). (B-D) Violin plots showing median and 25th and 75th percentile of (B) NAE capacity, (C) calculated titratable acidity (TA), and (D) acid-base (AB) score. (E-H) Pearson correlation analysis between urine pH and urine calcium (E), NAE and urine calcium (F), urine ammonium and calcium (G), and calculated TA and urine calcium (H). Blue dots = non stone formers (NSF), orange dots = CaOx stone formers, black dots = CaP stone formers. R = Pearson correlation coefficient and p = p-value associated with this correlation. α = 0.05.

Supplemental Figure 4: No major differences were found between COM and COD CaOx stone formers in relation to associations between acid-base parameters and calcium excretion. (A) 24h ammonium excretion. (B) Urine pH. (C) 24h citrate excretion (TA). (D) 24h urine calcium. (E) Pearson correlation analysis between urine pH and NAE. (F) NAE capacity (G). Pearson correlation analyses between urine pH and urine calcium (G), and NAE and urine calcium (H). Violin plots show median and 25th and 75th percentile. Blue dots = non stone formers (NSF), orange dots = COM stone formers, purple dots = COD stone formers, black dots = CaP stone formers. R = Pearson correlation coefficient and p = p-value associated with this correlation. α = 0.05.

