## Supplementary figures for "Acid excretion is impaired in calcium oxalate stone formers"

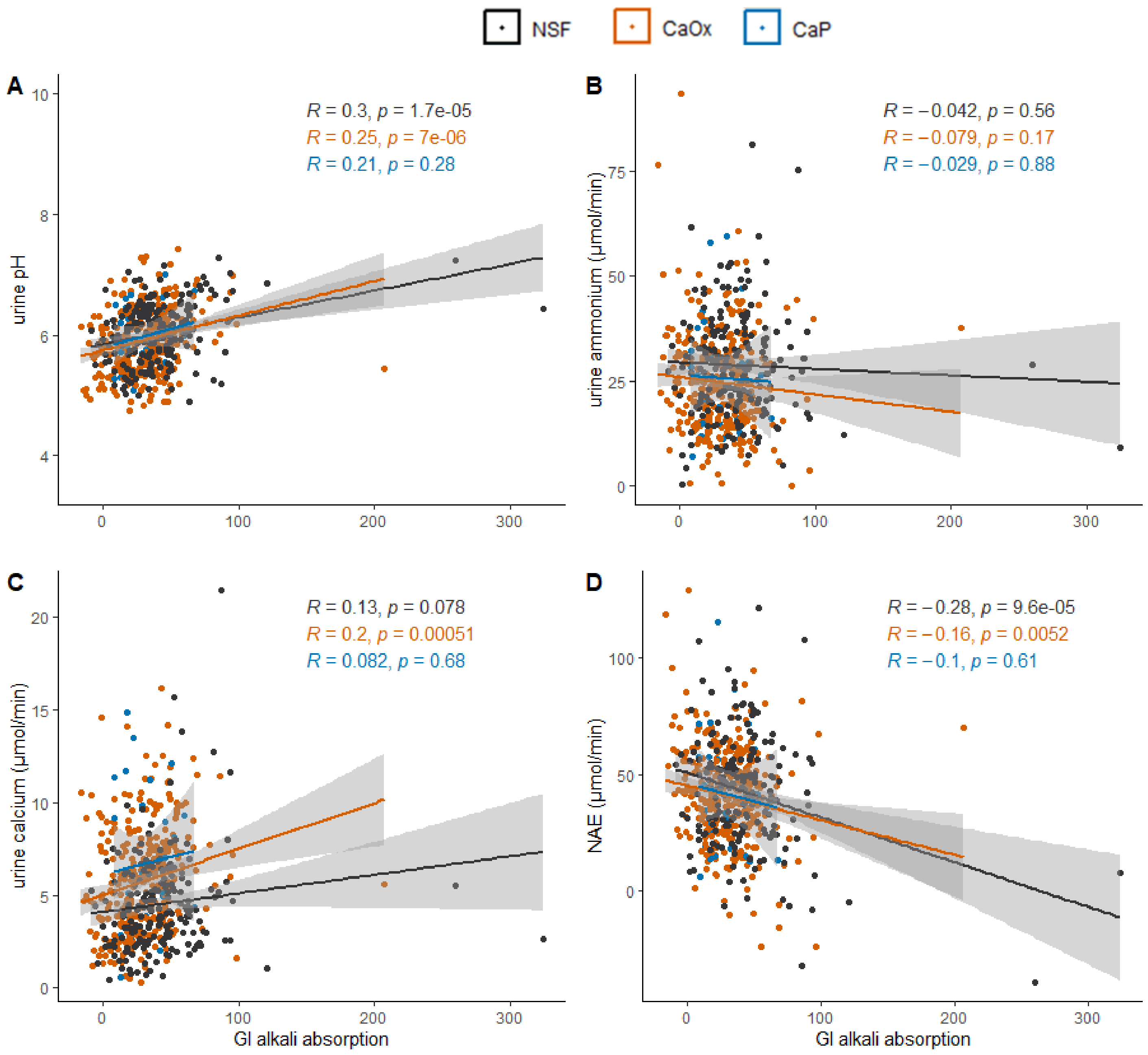


Supplemental Figure 1: Variation in gastrointestinal alkali absorption does not explain altered ammonium or calcium excretion. Correlation between calculated gastrointestinal alkali absorption and urine pH (A), ammonium (B), calcium (C), and net acid excretion is not significantly modified in CaOx or CaP stone formers. Black dots = NSF, orange dots = CaOx stone formers, and blue dots = CaP stone formers. R = Pearson correlation coefficient and p = p-value associated with this correlation. α = 0.05.

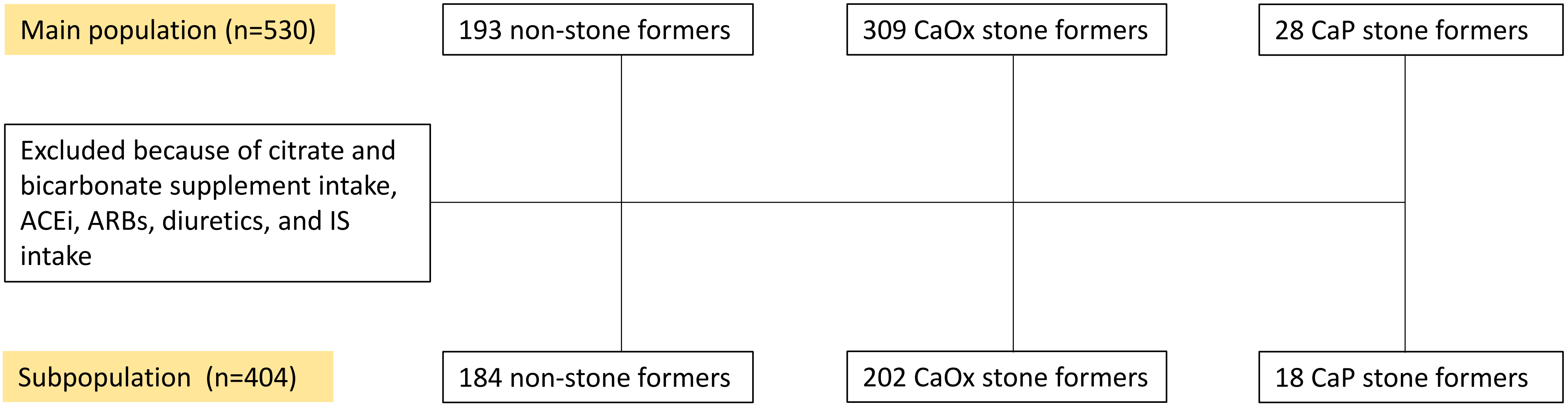


Supplemental Figure 2: Number of individuals per group after exclusion by drugs that affect the acid-base status. ACEi = Angiotensin-converting enzyme inhibitors, ARBs = Angiotensin receptor blockers, IS = immunosuppressants

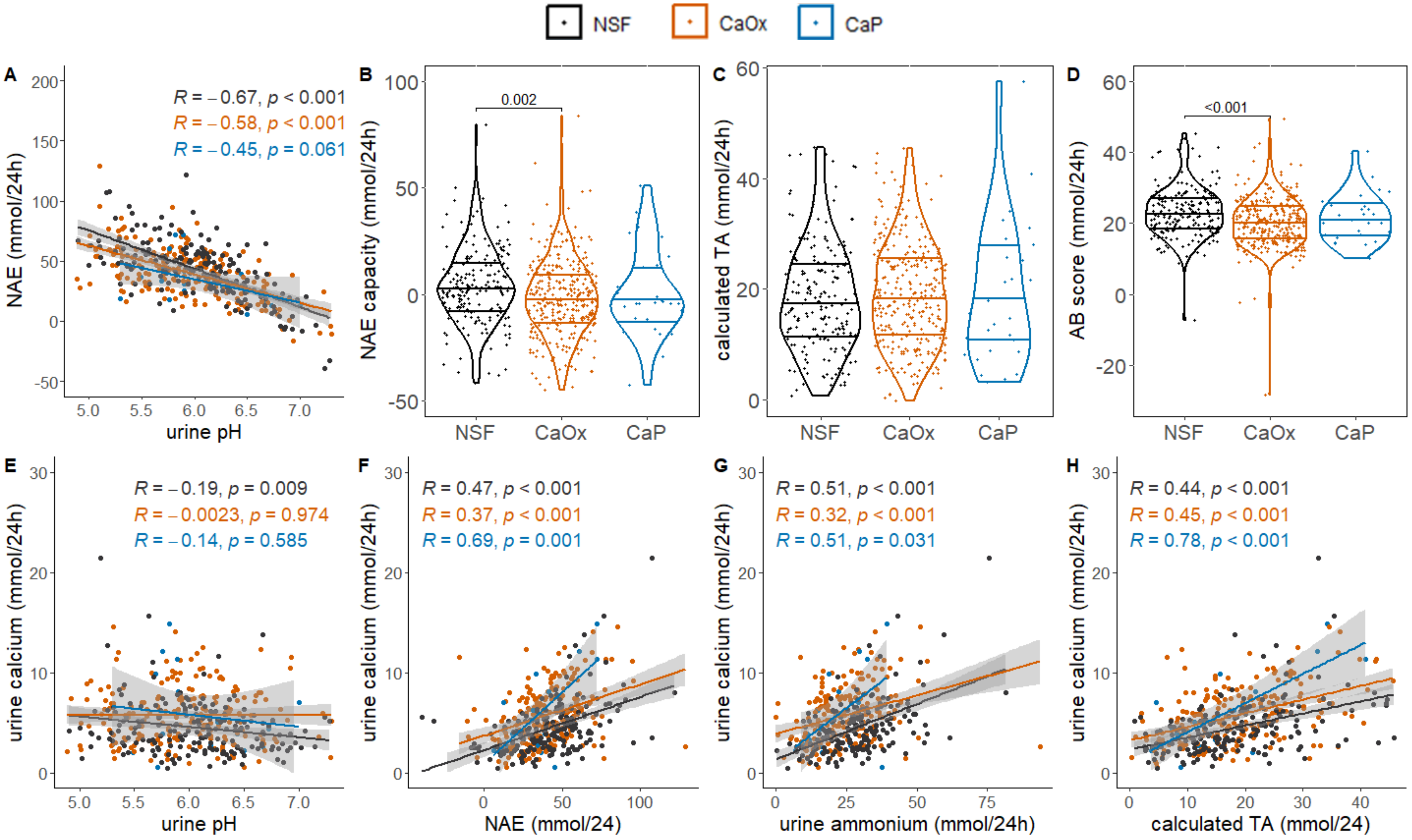


Supplemental Figure 3: Intake of medication and supplements interacting with acid-base balance did not play a major role in acid excretion capacity and its relationship with calcium excretion. (A) Pearson correlation analysis between urine pH and net acid excretion (NAE). (B) NAE capacity. (C) Calculated titratable acidity (TA). (D) Calculated acid-base (AB) score. (E-H) Pearson correlation analysis between urine pH and urine calcium (E), NAE and urine calcium (F), urine ammonium and calcium (G), and calculated TA and urine calcium (H). Blue dots = non stone formers (NSF), orange dots = CaOx stone formers (SF), black dots = CaP SF. R = Pearson correlation coefficient and p = p-value associated with this correlation. α = 0.05.
