## Supplementary tables for "Acid excretion is impaired in calcium oxalate stone formers"

| ATC Code | Common name of the medication or supplement |
|---|---|
| A12BA02 | Potassium citrate |
| A12BA04 | Potassium Bicarbonate |
| A12CC04 | Magnesium citrate |
| B05CB02 | Sodium citrate |
| B05CB04 | Sodium bicarbonate (irrigating solution) |
| B05XA02 | Sodium bicarbonate (solution additive) |
| A02BC02 | Pantoprazole |
| C03AA03 | Hydrochlorothiazide |
| C09AA01 | Captopril |
| C09AA02 | Enalapril |
| C09AA03 | Lisinopril |
| C09BA02 | Enalapril and diuretics |
| C09BA03 | Lisinopril and diuretics |
| C09BA05 | Ramipril + hydrochlorothiazide |
| C09CA06 | Candesartan |
| C09CA06 | Candesartan + hydrochlorothiazide |
| C09CA08 | Olmesartan medoxomil |
| C09DA01 | Losartan and diuretics |
| C09DA03 | Valsartan + hydrochlorothiazide |
| C09DA04 | Ibesartan + hydrochlorothiazide |
| C09DA08 | Olmesartan medoxomil and diuretics |
| C09DB02 | Olmesartan medoxomil and amlodipine |
| C09DX03 | Olmesartan medoxomil, amlodipine and hydrochlorothiazide |

| | |
|---|---|
| L04AD01 | Cyclosporine |
| S01EC01 | Acetazolamide |

Supplemental Table 1. Medication and supplements used as exclusion criteria in subpopulations 1. ATC code = The Anatomical Therapeutic Chemical code.

| Blood parameters | | | |
|---|---|---|---|
| | NSF (N=193) | CaOx (N=309) | CaP (N=28) |
| Creatinine (mmol/l) | 76.4 ± 13.5, n = 192 | 77.5 ± 17.6, n = 308 | 69.9 ± 17.9, n = 27 |
| pH | 7.38 ± 0.031, n = 71 | 7.40 ± 0.03, n = 68 | 7.41 ± 0.03, n = 7 |
| PaCO2 (mmHg) | 43.6 ± 6.45, n = 100 | 43.6 ± 6.32, n = 113 | 38.1 ± 7.58, n = 10 |
| Chloride (mmol/l) | 104 ± 2.47 | 103 ± 2.81, n = 307 | 103 ± 2.0 |
| Potassium (mmol/l) | 3.99 ± 0.266 | 4.09 ± 0.311 | 3.97 ± 0.356 |
| Sodium (mmol/l) | 141 ± 2.05 | 141 ± 2.14 | 141 ± 2.20 |
| Inorganic phosphate (mmol/l) | 1.01 ± 0.160 | 0.98 ± 0.18 | 1.03 ± 0.21 |
| Magnesium (mmol/l) | 0.81 ± 0.05 | 0.82 ± 0.23 | 0.80 ± 0.05 |
| PTH (ng/l) | 37.1 ± 15.6 | 42.1 ± 24.2 | 41.9 ± 21.4, n = 27 |
| Calcidiol (mmol/l) | 57.0 [40.0, 75.0] | 52.0 [34.2, 71.0] | 49.5 [29.8, 70.3] |
| Calcitriol (mmol/l) | 116 ± 33.1 | 118 ± 37.4 | 124 ± 45.0 |
| Ionized calcium (mmol/l) | 1.19 ± 0.12, = 99 | 1.21 ± 0.18, n = 118 | 1.19 ± 0.04, n = 10 |
| FGF23 (pg/ml) | 43.7 ± 13.5, n = 191 | 44.6 ± 25.6, n = 304 | 45.0 ± 15.4, n = 26 |
| Cholesterol (mmol/l) | 4.79 ± 1.01 | 4.82 ± 1.09 | 4.48 ± 0.81, n = 27 |
| HDL (mmol/l) | 1.42 ± 0.37 | 1.30 ± 0.37, n = 175 | 1.39 ± 0.34, n = 14 |
| LDL (mmol/l) | 3.15 ± 0.96 | 3.10 ± 1.05, n = 175 | 2.84 ± 0.89, n = 14 |

Supplemental Table 2: Blood parameters in stone former individuals and healthy NSFs. Continuous data are shown as mean ± SD and continuous skewed data as median [25th percentile; 75th percentile]. Parameters with missing data have their sample size (n) shown next to the mean or median value of each group. PTH = parathyroid hormone, LDL = low-density lipoprotein, HDL = high-density lipoprotein, FGF23 = fibroblast growth factor 23. Student t-test was used for continuous variables. α = 0.05.

CaOx adjusted

| Urine parameter | estimate | Std error | statistic | p-value | OR | 2.50% | 97.50% |
|---|---|---|---|---|---|---|---|
| (Intercept) | 0.22 | 0.20 | 1.120 | 0.263 | 1.24 | 0.85 | 1.83 |
| Ammonium | -0.89 | 0.17 | -5.378 | <0.001 | 0.41 | 0.29 | 0.56 |
| pH | -0.27 | 0.12 | -2.163 | 0.031 | 0.77 | 0.60 | 0.97 |
| Citrate | -0.56 | 0.12 | -4.509 | <0.001 | 0.57 | 0.44 | 0.72 |
| Phosphate | 0.03 | 0.17 | 0.204 | 0.839 | 1.03 | 0.75 | 1.43 |
| Calcium | 1.06 | 0.16 | 6.644 | <0.001 | 2.89 | 2.14 | 4.00 |
| Magnesium | -0.22 | 0.15 | -1.441 | 0.150 | 0.80 | 0.60 | 1.08 |
| Oxalate | -0.14 | 0.15 | -0.906 | 0.365 | 0.87 | 0.63 | 1.16 |
| Age | 0.04 | 0.12 | 0.303 | 0.762 | 1.04 | 0.82 | 1.32 |
| BMI | 0.47 | 0.12 | 3.844 | <0.001 | 1.61 | 1.27 | 2.06 |
| Sex (male) | 0.65 | 0.25 | 2.584 | 0.010 | 1.91 | 1.17 | 3.14 |

Supplemental Table 3. Summary of results of the logistic regression model 1 with CaOx stone formation as dependent variable. OR = odds ratio. The p-values were obtained from a Wald test with α = 0.05.

CaOx with NAEC adjusted

| Urine parameter | estimate | Std error | statistic | p-value | OR | 2.50% | 97.50% |
|---|---|---|---|---|---|---|---|

| | | | | | | | |
|---|---|---|---|---|---|---|---|
| (Intercept) | 0.19 | 0.19 | 0.999 | 0.318 | 1.21 | 0.83 | 1.76 |
| NAEC | -0.73 | 0.15 | -4.867 | <0.001 | 0.48 | 0.36 | 0.64 |
| Citrate | -0.51 | 0.12 | -4.274 | <0.001 | 0.60 | 0.48 | 0.76 |
| Calcium | 1.04 | 0.16 | 6.607 | <0.001 | 2.84 | 2.11 | 3.92 |
| Magnesium | -0.23 | 0.14 | -1.573 | 0.116 | 0.80 | 0.60 | 1.06 |
| Oxalate | -0.23 | 0.15 | -1.536 | 0.125 | 0.79 | 0.58 | 1.05 |
| Age | -0.01 | 0.12 | -0.108 | 0.914 | 0.99 | 0.78 | 1.25 |
| BMI | 0.48 | 0.12 | 3.927 | <0.001 | 1.61 | 1.28 | 2.06 |
| Sex (male) | 0.68 | 0.24 | 2.833 | 0.005 | 1.98 | 1.24 | 3.19 |

Supplemental Table 4. Summary of results of the logistic regression model 1 with CaOx stone formation as dependent variable and NAEC replacing ammonium, pH, and phosphate. OR = odds ratio. The p-values were obtained from a Wald test with α = 0.05.

CaOx with AB adjusted

| Urine parameter | estimate | Std error | statistic | p-value | OR | 2.50% | 97.50% |
|---|---|---|---|---|---|---|---|
| (Intercept) | 0.29 | 0.19 | 1.505 | 0.132 | 1.33 | 0.92 | 1.95 |
| AB score | -0.31 | 0.12 | -2.551 | 0.011 | 0.73 | 0.58 | 0.93 |
| Citrate | -0.49 | 0.12 | -4.127 | <0.001 | 0.61 | 0.48 | 0.77 |
| Phosphate | -0.26 | 0.15 | -1.725 | 0.085 | 0.77 | 0.57 | 1.04 |
| Calcium | 0.96 | 0.16 | 6.105 | <0.001 | 2.60 | 1.93 | 3.57 |
| Magnesium | -0.27 | 0.15 | -1.838 | 0.066 | 0.76 | 0.57 | 1.02 |
| Oxalate | -0.28 | 0.15 | -1.836 | 0.066 | 0.76 | 0.55 | 1.00 |
| Age | 0.09 | 0.12 | 0.824 | 0.410 | 1.10 | 0.88 | 1.38 |
| BMI | 0.43 | 0.12 | 3.621 | <0.001 | 1.54 | 1.23 | 1.96 |
| Sex (male) | 0.50 | 0.24 | 2.076 | 0.038 | 1.66 | 1.03 | 2.67 |

Supplemental Table 5. Summary of results of the logistic regression model 1 with CaOx stone formation as dependent variable and NAEC replacing ammonium, pH, and phosphate. OR = odds ratio. The p-values were obtained from a Wald test with α = 0.05.

Subpopulation after exclusion by supplementation and drug intake

CaOx unadjusted

| Urine parameter | estimate | Std error | statistic | p-value | OR | 2.50% | 97.50% |
|---|---|---|---|---|---|---|---|
| (Intercept) | 0.09 | 0.12 | 0.770 | 0.441 | 1.09 | 0.87 | 1.38 |
| Ammonium | -0.78 | 0.18 | -4.430 | <0.001 | 0.46 | 0.32 | 0.64 |

| | | | | | | | |
|---|---|---|---|---|---|---|---|
| pH | -0.24 | 0.12 | -1.908 | 0.056 | 0.79 | 0.62 | 1.00 |
| Citrate | -0.39 | 0.14 | -2.883 | 0.004 | 0.68 | 0.51 | 0.88 |
| Phosphate | 0.24 | 0.17 | 1.420 | 0.156 | 1.27 | 0.91 | 1.79 |
| Calcium | 1.02 | 0.17 | 5.929 | <0.001 | 2.76 | 2.01 | 3.93 |
| Magnesium | -0.29 | 0.17 | -1.691 | 0.091 | 0.75 | 0.53 | 1.04 |
| Oxalate | -0.58 | 0.19 | -3.042 | 0.002 | 0.56 | 0.38 | 0.81 |

Supplemental Table 6. Summary of results of the logistic regression model 0 with CaOx stone formation as dependent variable in subpopulation 1. OR = odds ratio. The p-values were obtained from a Wald test with α = 0.05.

CaOx with NAEC unadjusted

| Urine parameter | estimate | Std error | statistic | p-value | OR | 2.50% | 97.50% |
|---|---|---|---|---|---|---|---|
| (Intercept) | 0.10 | 0.12 | 0.838 | 0.402 | 1.10 | 0.88 | 1.38 |
| NAEC | -0.48 | 0.15 | -3.201 | 0.001 | 0.62 | 0.46 | 0.83 |
| Citrate | -0.33 | 0.13 | -2.532 | 0.011 | 0.72 | 0.56 | 0.92 |
| Calcium | 1.07 | 0.17 | 6.242 | <0.001 | 2.92 | 2.11 | 4.14 |
| Magnesium | -0.28 | 0.17 | -1.689 | 0.091 | 0.75 | 0.54 | 1.04 |
| Oxalate | -0.63 | 0.19 | -3.350 | 0.001 | 0.53 | 0.37 | 0.76 |

Supplemental Table 7. Summary of results of the logistic regression model 0 with CaOx stone formation as dependent variable and NAEC replacing ammonium, pH, and phosphate in subpopulation 1. OR = odds ratio. The p-values were obtained from a Wald test with α = 0.05.

CaOx adjusted

| Urine parameter | estimate | Std error | statistic | p-value | OR | 2.50% | 97.50% |
|---|---|---|---|---|---|---|---|

| | | | | | | | |
|---|---|---|---|---|---|---|---|
| (Intercept) | -0.41 | 0.21 | -1.925 | 0.054 | 1.24 | 0.85 | 1.83 |
| Ammonium | -0.83 | 0.18 | -4.520 | <0.001 | 0.41 | 0.29 | 0.56 |
| pH | -0.17 | 0.13 | -1.255 | 0.210 | 0.77 | 0.60 | 0.97 |
| Citrate | -0.38 | 0.14 | -2.713 | 0.007 | 0.57 | 0.44 | 0.72 |
| Phosphate | -0.01 | 0.18 | -0.064 | 0.949 | 1.03 | 0.75 | 1.43 |
| Calcium | 1.07 | 0.17 | 6.187 | <0.001 | 2.89 | 2.14 | 4.00 |
| Magnesium | -0.25 | 0.18 | -1.449 | 0.147 | 0.80 | 0.60 | 1.08 |
| Oxalate | -0.56 | 0.20 | -2.851 | 0.004 | 0.87 | 0.63 | 1.16 |
| Age | -0.11 | 0.13 | -0.849 | 0.396 | 1.04 | 0.82 | 1.32 |
| BMI | 0.46 | 0.14 | 3.356 | 0.001 | 1.61 | 1.27 | 2.06 |
| Sex (male) | 0.84 | 0.28 | 2.951 | 0.003 | 1.91 | 1.17 | 3.14 |

Supplemental Table 8. Summary of results of the logistic regression model 1 with CaOx stone formation as dependent variable in subpopulation 1. OR = odds ratio. The p-values were obtained from a Wald test with α = 0.05.

CaOx with NAEC adjusted

| Urine parameter | estimate | Std error | statistic | p-value | OR | 2.50% | 97.50% |
|---|---|---|---|---|---|---|---|
| (Intercept) | -0.45 | 0.21 | -2.135 | 0.033 | 1.21 | 0.83 | 1.76 |
| NAEC | -0.70 | 0.17 | -4.213 | <0.001 | 0.48 | 0.36 | 0.64 |
| Citrate | -0.33 | 0.13 | -2.471 | 0.013 | 0.60 | 0.48 | 0.76 |
| Calcium | 1.06 | 0.17 | 6.145 | <0.001 | 2.84 | 2.11 | 3.92 |
| Magnesium | -0.28 | 0.17 | -1.639 | 0.101 | 0.80 | 0.60 | 1.06 |
| Oxalate | -0.61 | 0.20 | -3.116 | 0.002 | 0.79 | 0.58 | 1.05 |

| | | | | | | | |
|---|---|---|---|---|---|---|---|
| Age | -0.15 | 0.13 | -1.179 | 0.239 | 0.99 | 0.78 | 1.25 |
| BMI | 0.46 | 0.13 | 3.471 | 0.001 | 1.61 | 1.28 | 2.06 |
| Sex (male) | 0.90 | 0.27 | 3.293 | 0.001 | 1.98 | 1.24 | 3.19 |

Supplemental Table 9. Summary of results of the logistic regression model 1 with CaOx stone formation as dependent variable and NAEC replacing ammonium, pH, and phosphate in subpopulation 1. OR = odds ratio. The p-values were obtained from a Wald test with α = 0.05.

CaOx with AB unadjusted

| Urine parameter | estimate | Std error | statistic | p-value | OR | 2.50% | 97.50% |
|---|---|---|---|---|---|---|---|
| (Intercept) | 0.09 | 0.12 | 0.822 | 0.411 | 1.10 | 0.88 | 1.38 |
| AB score | -0.28 | 0.13 | -2.245 | 0.025 | 0.75 | 0.59 | 0.96 |
| Citrate | -0.31 | 0.13 | -2.409 | 0.016 | 0.73 | 0.57 | 0.94 |
| Phosphate | -0.02 | 0.15 | -0.120 | 0.904 | 0.98 | 0.73 | 1.33 |
| Calcium | 0.96 | 0.17 | 5.560 | 0.000 | 2.60 | 1.88 | 3.70 |
| Magnesium | -0.37 | 0.17 | -2.177 | 0.030 | 0.69 | 0.50 | 0.96 |
| Oxalate | -0.68 | 0.19 | -3.572 | 0.000 | 0.51 | 0.34 | 0.73 |

Supplemental Table 10. Summary of results of the logistic regression model 0 with CaOx stone formation as dependent variable and AB score replacing ammonium and pH in subpopulation 1. OR = odds ratio. The p-values were obtained from a Wald test with α = 0.05.

CaOx with AB adjusted

| Urine parameter | estimate | Std error | statistic | p-value | OR | 2.50% | 97.50% |
|---|---|---|---|---|---|---|---|
| (Intercept) | -0.36 | 0.21 | -1.682 | 0.093 | 0.70 | 0.46 | 1.06 |
| AB score | -0.21 | 0.13 | -1.585 | 0.113 | 0.81 | 0.62 | 1.05 |

| | | | | | | | |
|---|---|---|---|---|---|---|---|
| Citrate | -0.30 | 0.13 | -2.274 | 0.023 | 0.74 | 0.56 | 0.95 |
| Phosphate | -0.28 | 0.17 | -1.678 | 0.093 | 0.75 | 0.54 | 1.05 |
| Calcium | 0.98 | 0.18 | 5.600 | <0.001 | 2.67 | 1.92 | 3.81 |
| Magnesium | -0.33 | 0.17 | -1.944 | 0.052 | 0.72 | 0.51 | 1.00 |
| Oxalate | -0.68 | 0.20 | -3.426 | 0.001 | 0.51 | 0.34 | 0.74 |
| Age | -0.07 | 0.13 | -0.549 | 0.583 | 0.93 | 0.72 | 1.20 |
| BMI | 0.41 | 0.13 | 3.113 | 0.002 | 1.50 | 1.17 | 1.96 |
| Sex (male) | 0.74 | 0.28 | 2.670 | 0.008 | 2.09 | 1.22 | 3.60 |

Supplemental Table 11. Summary of results of the logistic regression model 1 with CaOx stone formation as dependent variable and AB score replacing ammonium and pH in subpopulation 1. OR = odds ratio. The p-values were obtained from a Wald test with α = 0.05.
